# Genome-wide CRISPR screens identified *C18orf32* as a novel regulator of lipid metabolism that mediates PFOA-induced toxicity

**DOI:** 10.64898/2025.12.23.696201

**Authors:** Chanhee Kim, Abderrahmane Tagmount, Zhaohan Zhu, W. Brad Barbazuk, Rhonda Bacher, Christopher D. Vulpe

## Abstract

Perfluorooctanoic acid (PFOA) remains a public health concern due to its persistence in the environment, despite global production bans. Epidemiological, animal, and *in vitro* studies have consistently linked PFOA exposure to hepatotoxicity characterized by lipid dysregulation; however, the molecular mechanisms underlying these adverse effects in humans remain poorly understood. To address this gap, we employed an integrated functional toxicogenomics framework combining CRISPR screening (for gene target identification), target validation, and assessment of target-derived cellular and molecular phenotypes. From genome-wide CRISPR screens in HepG2/C3A human liver cells, we identified 319 candidate genes (140 sensitive and 179 resistant) that modulate PFOA toxicity. Among these, *C18orf32*—encoding a lipid droplet (LD)-associated protein and the top resistant candidate—was selected for further mechanistic investigation based on its potential functional involvement in lipid metabolism. Targeted knockout (KO) of *C18orf32* conferred marked cellular resistance to PFOA, accompanied by reduced LD and Triglyceride accumulation in cells under the exposure condition, suggesting a functional role of C18orf32p in lipid dysregulation. To elucidate molecular mechanisms underlying the cellular phenotypes, we conducted systematic transcriptomics profiling of wild-type (WT) and *C18orf32* KO cells in the absence and presence of PFOA exposure. *C18orf32* KO caused extensive gene expression reprogramming, with over one-third of coding genes differentially expressed. Notably, a range of lipid metabolism pathways including cholesterol metabolism, PPAR signaling, fatty acid metabolism, and peroxisome β-oxidation—known mediators of PFOA-induced hepatotoxicity—were significantly downregulated in *C18orf32* KO cells, contrasting to upregulation of these pathways in PFOA exposed WT cells. Collectively, our results identify C18orf32p as a previously unrecognized regulator of hepatic lipid metabolism and a key genetic determinant of PFOA-induced lipid dysregulation and hepatotoxicity in humans.

## Introduction

Per– and polyfluoroalkyl substances (PFAS) are a large class of synthetic chemicals widely used in industrial and consumer products for their surfactant and non-stick properties. Among them, perfluorooctanoic acid (PFOA), as a legacy PFAS, has been extensively studied due to its persistence, bioaccumulation, and adverse health effects in both humans and wildlife^1–4^. Despite the global phaseout of PFOA production, its environmental persistence and continued detection in human serum underscore the need for a mechanistic understanding of PFOA-induced adverse outcomes to provide guidance on effective risk assessment and inform preventive and therapeutic strategies to mitigate PFOA exposure^5–8^. While many adverse health effects are reported for PFOA, such as reproductive toxicity, endocrine toxicity, and immunotoxicity^4,9,10^, mounting toxicological and epidemiological evidence links chronic PFOA exposure to metabolic disturbances, hepatomegaly, steatosis, elevated liver enzymes in serum, and increased risk of liver disease, suggesting its hepatotoxic potential in humans^11–14^. Experimental studies in rodents and human *in vitro* models have also documented that PFOA exposure induces hepatocyte hypertrophy, lipid accumulation, and metabolic dysfunction^14–17^. However, despite decades of extensive research, the molecular mechanisms underlying PFOA-induced liver toxicity remain incompletely defined, particularly regarding how PFOA perturbs lipid metabolism and homeostasis^16,18^.

Peroxisome proliferator-activated receptor alpha (PPARα) activation does not fully account for PFOA hepatotoxicity in people. Rodent studies demonstrate that PFOA can function as a potent PPARα agonist resulting in peroxisome proliferation, hepatomegaly, and alterations in lipid metabolism^16,19,20^ but PFOA-induced outcomes are highly strain– and diet-dependent ^20,21^. In contrast, human hepatocytes exhibits markedly reduced responsiveness to PFOA and other PPARα agonists^22–25^. Emerging evidence support PFOA mediated disruption of mitochondrial function, endoplasmic reticulum (ER) stress^26,27^, and oxidative injury^28,29^, independent of PPARα signaling. Further, PFOA exposure has been linked to lipid droplet^16,30^. (LDs) accumulation in *C. elegans*^31^, mouse hepatocytes^32^, bovine blastocytes (exposed to PFNA, one carbon chain longer than PFOA)^33^, as well as human hepatic cells (HepaRG)^34,35^. Other signaling pathways including PPARγ, PPARδ, constitutive androstane receptor (CAR), pregnane X receptor (PXR), farnesoid X receptor (FXR), and retinoid X receptor (RXR)^25,36,37^which collectively regulate lipid storage, bile acid synthesis, glucose metabolism, and detoxification pathways^16,38–41^ have been proposed to play a role in PFOA mediated toxicity in people. Together, these findings underscore the continuing uncertainty about PFOA’s molecular targets and highlight the need for unbiased, systematic approaches to identify functional mediators that causally shape cellular responses to PFOA exposure.

Toxicogenomics approaches have provided valuable insights into PFOA exposure, revealing dysregulation of lipid metabolism, xenobiotic processing, liver diseases-related pathways, and inflammatory signaling^14,16,17,39,42,43^. However, distinguishing direct effects from secondary or downstream responses are challenging with these approaches. CRISPR-based functional genomics^44^ enables systematic loss-of-function interrogation of the entire genome to identify gene products that modulate chemical-induced toxicity^45–49^. Recent applications of genome-scale CRISPR screens have uncovered previously unrecognized key functional determinants of chemical and xenobiotic responses^49–51^, providing a causal framework for mechanistic toxicology. Despite this promise, the application of CRISPR screens to emerging chemicals of concern, particularly PFAS, remains limited^52^.

In this study, we conducted a genome-wide CRISPR/Cas9 knockout screen of PFOA toxicity in HepG2/C3A human hepatoma cells to identify candidate genes whose loss significantly modulate PFOA-induced cytotoxicity. In particular, *C18orf32,* which encodes a lipid droplet (LD)-associated protein not previously linked to PFOA toxicity, was identified as key candidate which significantly increased PFOA resistance when disrupted. We further validated the PFOA resistance in independent *C18orf32* KO cell lines and further characterized the functional role of C18orf32p in PFOA-induced lipid metabolism dysregulation. This work supports a key role C18orf32p in PFOA-driven hepatic lipid metabolic dysregulation, providing mechanistic insight into the molecular basis of PFOA-mediated liver toxicity.

## Materials and Methods

### 1. Cell cultures and PFOA cytotoxicity

HEK293T (for lentivirus production) and HepG2/C3A cells were purchased from the American Type Culture Collection (ATCC, Manassas, Virginia). Cells were maintained and passaged following ATCC’s recommended protocol. The HEK293T cells were cultured in Dulbecco’s Modified Eagle Media (DMEM; Thermo Fisher Scientific, Waltham, MA) and the HepG2/C3A cells were cultured in Minimum Essential Medium (MEM, Gibco). The complete media were supplemented with 10 % fetal bovine serum (FBS, Thermo Fisher Scientific, Waltham, MA), 1X MEM Non-Essential Amino Acids Solution 100X (Gibco, added for HepG2/C3A) and 1X Antibiotic-Antimycotic 100X (Gibco). Cells were cultured in a humidified incubator (Forma™ Series II Water-Jacketed CO2 Incubator, Thermo Scientific™) with 5% CO_2_ at 37 °C. Cytostasis/Cytotoxicity of PFOA (purity≥95%, Santa Cruz Biotechnology-sc250662, CAS# 335-67-1) exposure in HepG2/C3A cells exposed to a range of nominal concentrations (0–400 µM) in triplicate for 6 days was assessed by measuring ATP levels using the CellTiter-Glo2.0 cell viability assay kit (Promega, Madison, WI) following the manufacturer’s instruction and our previous study. Briefly, 100 µl of the CellTiter-Glo2.0 reagent (the same volume of the culture media) was added to each well containing cells and media and the plate was incubated for 15 min at room temperature in the dark to produce and stabilize the luminescence signal (relative light unit, RLU). The signals were read on a Synergy H1 microplate reader (BioTek Instruments, Winooski, VT). Relative luminescence signals were calculated and subsequently converted to % of viable cells compared to Dimethyl sulfoxide (DMSO) controls (PFOA concentration: 0 µM). The % values were used to determine 25 % inhibitory concentrations (IC_25_) for PFOA using GraphPad Prism (version 10.1.0) dose-response sigmoid function. Of note, we chose the duration of exposure as 6 days for the 25 % inhibitory concentration (IC_25_) calculation since PFOA exposure didn’t show detectable cytotoxicity in HepG2/C3A cells after 3 days of exposure, which is the exposure period that we routinely use to estimate sublethal concentrations for CRISPR screens.

### 2. Lentiviral production and transduction

HEK293T cells were used to produce lentivirus by co-transfection of the MibLibCas9 library plasmids, the packaging plasmid psPAX2 (#12260, Addgene), and envelope plasmid pMD2.G (#12259, Addgene). The MinLibCas9 lentiviral library was functionally tittered in HepG2/C3A cells to determine the amount of virus required to obtain a multiplicity of infection of 0.3-0.5. For large-scale transduction for the screens, HepG2/C3A cells were seeded in four 12-well culture plates (1×10^6^ cells/well). After a 24 h incubation period, polybrene (Sigma) was added at a concentration of 8 µg/ml along with 0.5 µl (Multiplicity of infection, MOI= 0.3) of the titrated MinLibCas9 lentiviral library in each well, followed by centrifugation (spinfection) at 33 ◦C at 1000xG for 2 h. After further incubation at 37 ◦C for 30 h, the lentiviral mix was replaced with 1 mL of the growth media in each well. After 48 h recovery period, the pooled cells were treated with puromycin (2 μg/mL) for 5 days to enrich for transduced cells before chemical exposures, resulting in almost no cells in the non-transduced control flask.

### 3. Genome-wide CRISPR loss-of-function screening

For the genome-wide loss-of-function genetic screens, we used a customized minimal genome-wide sgRNA library termed MinLibCas9 based on a published study which selected two sgRNAs with maximal on-target activity for each gene based on computational analysis of the screening results of several genome-wide screens^53^. The sgRNA library includes a total of 37,843 sgRNAs (two sgRNAs/gene, targeting total of 18819 genes, 200 non-targeting control sgRNAs, and 5 safe-harbor sites control sgRNAs) (Table S1). Lentivirus production was carried out as previously described^46^. A pool of the MinLibCas9 lentiviral library was transduced into HepG2/C3A cells at 0.3 multiplicity of infection (MOI) to generate a mutant pool as in previous studies^49,54^. The mutant pool was then expanded for the PFOA exposure experiments to obtain sufficient number of cells for replicates and controls. Control and PFOA exposure samples of 16 × 10^6^ cells in T225 flasks (three replicates) were used for screening. Exposure samples were treated with PFOA at inhibitory concentration (IC_25_) for 18 days (10 doubling times). Representation of the library was maintained throughout the screening experiment at approximately 400X the MinLibCas9 library size for each replicate.

### 4. DNA extraction, library preparation, and next-generation sequencing (NGS)

Genomic DNA was extracted from 1.6 × 10^7^ cells of each sample using the Quick-DNA Midiprep Plus Kit (ZYMO Research) following the manufacturer’s protocol. Amplicons for NGS Illumina sequencing were generated using the pairs of universal CRISPR-FOR1 forward primer and CRISPR-REV**#** reverse primers (**#**: 1 to 48) specific for each sample (see Table S2). Amplicons amplifying the sgRNA region in each sample were then pooled and gel purified using the QIAquick Gel Extraction Kit (Qiagen) and quantified using the Qubit HS dsDNA assay (Thermo Scientific). Equimolar amounts of each amplicon library were multiplexed in one pool. The NGS was carried out at the Interdisciplinary Center for Biotechnology Research (ICBR), University of Florida at Gainesville, using the NovaSeqX paired 150 bp high-throughput platform (Illumina).

### 5. Data processing and bioinformatics of CRISPR screening

After data acquisition (raw fastq.gz files), read quality was checked using FASTQC tools. A Model-based Analysis of Genome-wide CRISPR-Cas9 Knockout (MaGeCK)^55^ was then used to demultiplex raw fastq data files, which were further processed to generate reads containing only the unique 20 bp guide sequences. The resulting read counts were mapped to sgRNAs, and the counts were then normalized to adjust for the effect of library sizes and read count distributions. The sgRNAs were then ranked by a modified robust ranking aggregation (α-RRA) algorithm to generate a ranked gene list. Candidate genes were then defined as genes with *p*-value < 0.01 and Log_2_FC < −0.6 (sensitive) or Log_2_FC > 0.6 (resistant) for pathway analysis (GO-BP, KEGG, and MSigDB). We define sensitive genes as those that when disrupted by CRISPR targeting, show increased cellular sensitivity (less abundant than control) to PFOA exposure whereas resistant genes show increased resistance (more abundant than control) to PFOA exposure. We cross-validated the MaGeCK-produced candidate gene hit with the result obtained using Platform-independent Analysis of Pooled Screens using Python (PinAPL-Py)^56^.

### 6. Generation of a clonal *C18orf32* knockout cell line

We performed single-gene CRISPR/Cas9 biallelic targeted disruption of *C18orf32* in HepG2/C3A hepatic cells using sgRNA#1 targeting *C18orf32* (Table S1) selected from the MinLibCas9 sgRNA library. The sgRNA#1 was cloned into the LentiCRISPRv2 using the golden gate assembly strategy as described elsewhere^44^. Following generation of a *C18orf32* knockout (KO) mutant pool, we isolated and expanded multiple single-cell–derived clones which were screened by PCR of targeted region (Forward primer: CAAGGTCTCACTTTGTTGTG; Reverse primer: CAGCATAATCTCTCCTCATCC) followed by Sanger-sequencing (GENEWIZ). The resulting abi. files (chromatograms) were used to run DECODR^57^ analysis to identify biallelic predicted disruption. We further characterized two *C18orf32* KO clones (clone#1 and clone#2) by quantitative qPCR to assess transcript levels. Total RNA was isolated from wild-type (WT) and *C18orf32* knockout (KO) cells using the RNeasy Plus Mini Kit (Qiagen, Cat# 74134) according to the manufacturer’s protocol, including on-column DNase I digestion to remove genomic DNA contamination. RNA concentration and purity were assessed by a NanoDrop spectrophotometer (Thermo Fisher Scientific), and 500 ng of total RNA was reverse-transcribed into cDNA using an iScript^TM^ cDNA synthesis Kit (Bio-Rad). Quantitative PCR (qPCR) was performed using SYBR Green master mix (Bio-Rad) on a real-time PCR system under standard cycling conditions. Gene-specific qPCR primers (qPCR_F: CCTCTGGTTTCCCCCTTCGTTA, qPCR_R: CACAGATTTCTGTTGGTCCTTTTG) were used to amplify *C18orf32*, and β-actin (*ACTB*) was used as an internal reference gene. Relative mRNA expression levels were calculated using the 2⁻ΔΔCt method^58^, with WT samples serving as the calibrator.

### 7. Fluorescence cellular imaging of lipid droplets

Cellular neutral lipid content (lipid droplet) was assessed using the fluorescent lipophilic dye BODIPY 493/503 (4,4-difluoro-1,3,5,7,8-pentamethyl-4-bora-3a,4a-diaza-s-indacene), which selectively stains triglycerides and cholesterol esters within lipid droplets. Briefly, cells were seeded into 96-well culture vessels and treated as indicated. Following treatment, cells were washed twice with phosphate-buffered saline (PBS) and incubated with BODIPY 493/503 (1 ug/ml in PBS) for 30 min in the dark to label intracellular neutral lipids. Nuclei were counterstained with NucBlue (Hoechst 33342, 2 drops/ml) simultaneously. Fluorescent images were acquired using a KEYENCE BZ-X800 fluorescence microscope equipped with appropriate filters (excitation/emission: 493/503 nm for BODIPY). Exposure and acquisition settings were kept constant across all experimental conditions.

### 8. Cellular Triacylglycerol (TAG) measurement

Cellular triacylglycerol (TAG) levels are quantified using a Triglyceride-Glo^TM^ Assay (Promega) following the manufacturer’s instructions with minor modifications. Briefly, WT and *C18orf32* KO cells were seeded in 96-well plates and exposed to PFOA at 100 μM for 72 h. Luminescence was measured using a Synergy H1 microplate reader (BioTek) with standard settings. TAG content was normalized to total cell number, and all experiments were performed triplicate.

### 9. Transcriptome profiling (RNA-seq)

#### RNA isolation and Quality assessment

HepG2/C3A cell samples representing four experimental conditions in triplicate: wild-type (WT), *C18orf32* knockout (KO), WT treated with PFOA (100 μM, 72 h), and KO treated with PFOA (100 μM, 72 h), were harvested at ∼ 80 % confluency for whole-transcriptome profiling. Total RNA was isolated using the RNeasy Plus Mini Kit (Qiagen, Cat# 74134) according to the manufacturer’s protocol, including on-column DNase I digestion to remove genomic DNA contamination. RNA concentration and purity were measured using a NanoDrop spectrophotometer (Thermo Fisher Scientific), and RNA integrity was assessed using the Agilent 4200 TapeStation (Agilent Technologies). Samples with RNA integrity number (RIN) > 7.5 and distinct 18S/28S peaks were deemed suitable for library preparation and all samples met the criteria. A minimum of 500 ng total RNA per sample was used for library preparation.

#### Library preparation and RNA-seq

RNA samples were submitted to the University of Florida Interdisciplinary Center for Biotechnology Research (UF ICBR)-Gene expression and Genotyping Core for RNA-seq services. Library preparation was performed by UF ICBR using poly(A) selection to enrich for mRNA, followed by strand-specific cDNA synthesis. Sequencing libraries were constructed using the Illumina TruSeq Stranded mRNA Library Prep Kit according to the standard ICBR workflow. Prepared libraries were quantified, pooled, and sequenced on the Illumina (NovaSeqX) platform to generate 150-bp paired-end reads. Sequencing depth targeted 30-40 million read pairs per sample. RNA-seq of 12 samples (triplicates per group) generated an average of 51 million paired-end reads per sample, with > 95 % uniquely mapped to the human genome (GRCh38) ([iDEP](https://bioinformatics.sdstate.edu/idep/))

#### Preprocessing and Quality control

Raw sequencing reads were processed by UF ICBR for initial quality filtration, including removal of adaptor sequence and low-quality bases. FastQC metrics were reviewed to ensure sufficient base quality scores and sequence complexity. Post-delivery, an additional in-house quality check was performed using PartekFlow (Partek Inc.). Reads were then aligned to the human reference genome (GRCh38) using the STAR aligner under default PartekFlow parameters. Gene-level quantification was performed using PartekFlow’s implementation of the Expectation/Maximization algorithm based on GENCODE_v47 annotations.

#### Differential Expression Analysis (DESeq2) and Pathway Enrichment analysis

Downstream expression analysis was performed in PartekFlow. Read counts were normalized using the median ratio method and differentially expressed genes (DEGs) were identified using DESeq2 module implemented by PartekFlow. Primary comparisons were made between (significance thresholds per comparison): 1. *C18orf32* KO vs. WT (FDR < 0.01 & |Log_2_FC| >2), 2. WT +PFOA vs. WT (FDR < 0.05 & |Log_2_FC| >0.5), *C18orf32* KO +PFOA vs. *C18orf32* KO (FDR < 0.05 & |Log_2_FC| >0.5), and 3. *C18orf32*

KO +PFOA vs. WT (FDR < 0.01 & |Log_2_FC| >2). The resulting lists of DEGs from each comparison were used for functional enrichment analysis. GO-BP, KEGG, MSigDB functional enrichment analyses was conducted using ShinyGO 0.85.1 ([ShinyGO 0.85](https://bioinformatics.sdstate.edu/go/))^59^ and the two-directional plot was generated using clusterProfiler implemented by SRPLOT software^60^.

## Results

### 1. CRISPR screens identify *C18orf32* as the top candidate gene modulating PFOA toxicity

To identify genes that modulate PFOA-induced cellular toxicity, genome-wide CRISPR loss-of-function screens were performed in HepG2/C3A cells exposed to a sublethal PFOA dose as illustrated in Fig. 1. We first determined IC_25_ dose of PFOA, which inhibits cellular proliferation by 25 % compared to no exposure control for 6 days exposure, to be 125 µM (Supplementary Figure S1). We used exposure to the IC_25_ of PFOA in an effort to identify genes which confer sensitivity or resistance to PFOA, when genetically disrupted^45,61^. Using the MinLibCas9 sgRNA library (37,843 sgRNAs, two sgRNAs per gene), a mutant cell pool was generated and either maintained as a control or exposed to PFOA (IC_25_= 125 µM) for 18 days (approximately 10 doublings). Candidate genes affecting PFOA sensitivity or resistance were identified by comparing relative sgRNA abundance between control and PFOA-exposed cells determined by next generation sequencing (NGS) and MaGeCK (α-RRA) gene ranking analysis^55^, with thresholds of p-value < 0.01 and log_2_FC < –0.6 (sensitive) / > 0.6 (resistant) (Figure 1B). This analysis identified a total of 319 candidate genes (140 sensitive genes and 179 resistant genes) that, when disrupted, modulate PFOA toxicity (Figure 1C, Table S3). Functional enrichment analyses of these sensitive or resistant candidate genes showed no significant enrichment with GO-BP and KEGG databases but showed enrichment with MSigDB (v7.5) Hallmark gene sets (Supplementary Figure S2). The PFOA-sensitive genes were enriched for several lipid metabolism-related terms such as fatty acid metabolism, oxidative phosphorylation, bile acid metabolism, and peroxisome and immune response-related terms such as interferon gamma/alpha response (Supplementary Figure S2A). Other enriched terms include ROS pathway, Hypoxia, mTORC1signaling, and G2-M checkpoint. The PFOA-resistant genes were also enriched for several lipid metabolism-related terms such as peroxisome and cholesterol homeostasis and reproductive terms such as spermatogenesis and mitotic spindle, and others include glycolysis, DNA repair, xenobiotic metabolism, and unfolded protein response (Supplementary Figure S2B).

**Figure 1.**
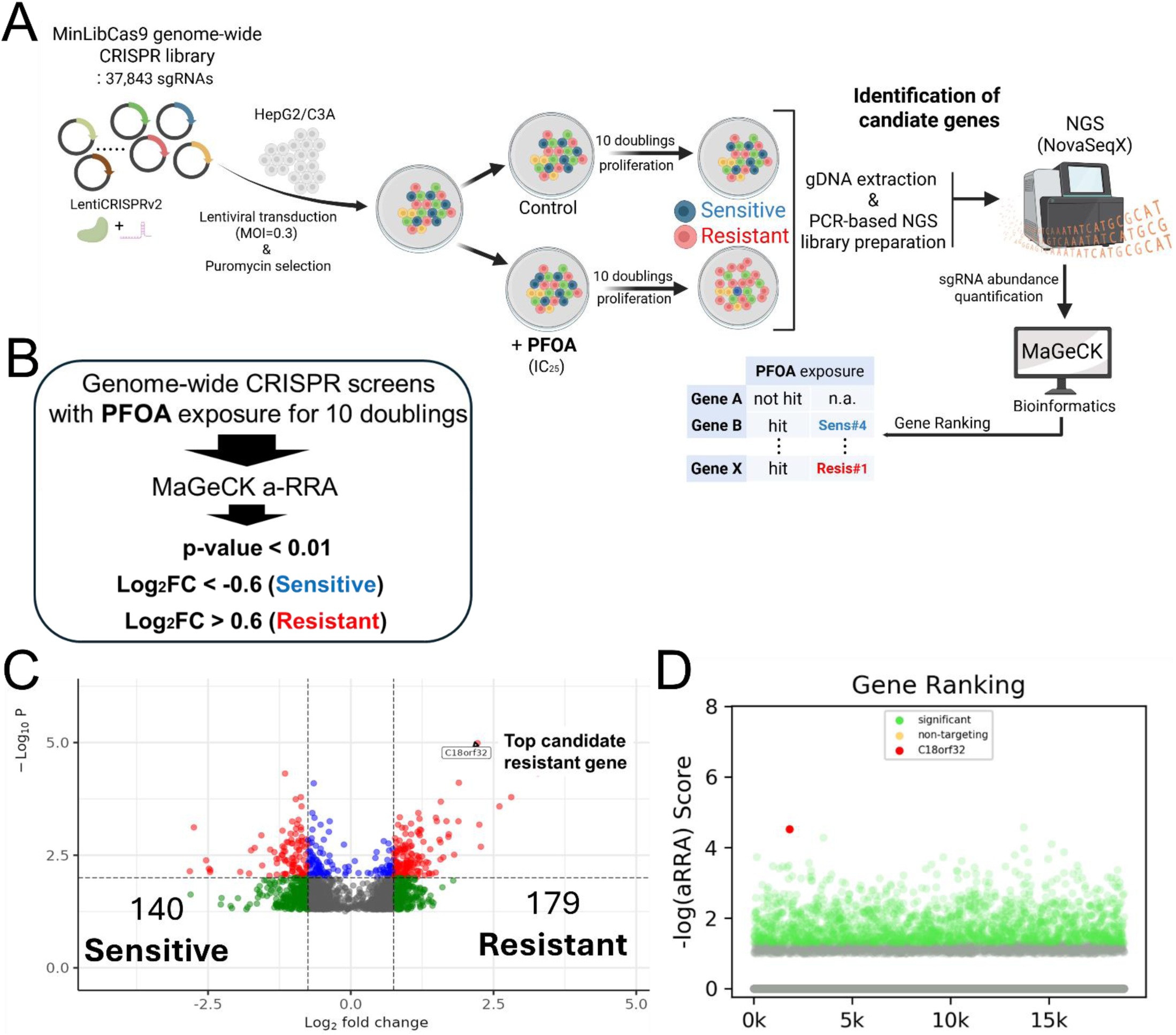
Genome-wide CRISPR knockout screen identifies *C18orf32* as a top genetic determinant of PFOA-induced cytotoxicity. (A) Schematic overview of the genome-wide CRISPR/Cas9 loss-of-function screen. HepG2/C3A cells stably expressing Cas9 were transduced with the MiniLibCas9 sgRNA library (37,843 sgRNAs; MOI = 0.3) and selected with puromycin. Cells were then split into control and PFOA-exposed groups and cultured for ∼10 doublings. PFOA treatment (IC₅ concentration) selectively depleted sgRNAs from sensitive gene knockouts and enriched sgRNAs from resistant gene knockouts. Genomic DNA was isolated, sgRNA cassettes were PCR-amplified, and next-generation sequencing (NovaSeqX) was performed. sgRNA abundance was quantified and analyzed using MAGeCK to rank gene-level enrichment or depletion. MOI, Multiplicity of Infection (MOI); IC_25_, Inhibitory Concentration of 25 %. (B) Schematic depiction of bioinformatics flow to identify candidates in the screen: sensitive genes (blue), whose loss increases PFOA toxicity, and resistant genes (red), whose loss confers resistance to PFOA. (C) Volcano plot showing gene-level log₂ (fold-change) versus –log(p-value) from MAGeCK analysis. A total of 140 sensitive genes and 179 resistant genes were significantly altered under PFOA exposure. *C18orf32* is highlighted as the top candidate resistant gene. (D) Ranked −log(αRRA) resistant gene scores from the PinAPL-Py analysis. Each point represents one gene in the CRISPR library. *C18orf32* (red) ranks among the most significant resistant hits, well above the distribution of background or non-targeting control guides.

Among the 319 candidate genes, *C18orf32* was selected for downstream mechanistic investigation because it emerged as the top PFOA-resistant hit in our screen (Figure 1C) and encodes a recently characterized lipid droplet (LD)-associated protein^62^ potentially relevant to PFOA-mediated lipid dysregulation. This choice was further supported by our functional enrichment analyses, which revealed that both PFOA-sensitive and PFOA-resistant gene sets were significantly enriched for lipid metabolism pathways, suggesting that lipid regulatory networks play a central role in modulating PFOA toxicity. Also, in addition to the MaGeCK (α-RRA) analysis, we independently validated *C18orf32* as a top candidate using PinAPL-Py^56^ analysis, which also ranked it among the strongest candidate hits (Figure 1D). Together, these results highlighted *C18orf32* as a high-confidence genetic modulator of PFOA toxicity and a biologically relevant gene for further mechanistic characterization in this study.

### 2. Targeted *C18orf32* knockout validates the phenotype observed in the screens

The clonal *C18orf32* KO cell lines (clone#1 and clone#2) generated by CRISPR/Cas9 using sgRNA#1 (Figure 2A and Supplementary Figure S3) were predicted to harbor 100% frameshift-inducing INDELs, with *C18orf32* KO_clone#1 carrying –41 (52.1%) and –20 (47.9%) bp deletions, and KO_clone#2 carrying a uniform –7 bp deletion (Figure 2B). Consistent with these mutations, both clones exhibited near-complete loss of *C18orf32* mRNA expression relative to wild-type (WT) cells (relative normalized expression: 0.0008 for clone #1 and 0.02 for clone #2), confirming effective gene disruption (Figure 2C). Functionally, both *C18orf32* KO clones showed significant cellular resistance to PFOA exposure over a 6-day period, with improved viability compared to WT cells. Clone #1 displayed a more pronounced resistance phenotype at 150 µM PFOA (2.7-fold increase compared to WT, p= 0.00028) relative to clone #2 (1.4-fold increase compared to WT, p= 0.03) (Figure 2D). Shorter exposure assays (2 and 4 days) further confirmed consistent and reproducible resistance phenotypes in both KO clones (Figure 2E and 2F). Together, these results demonstrate that targeted disruption of *C18orf32* recapitulates the resistance phenotype observed in our CRISPR screens, with clone #1 exhibiting the strongest functional effect.

**Figure 2.**
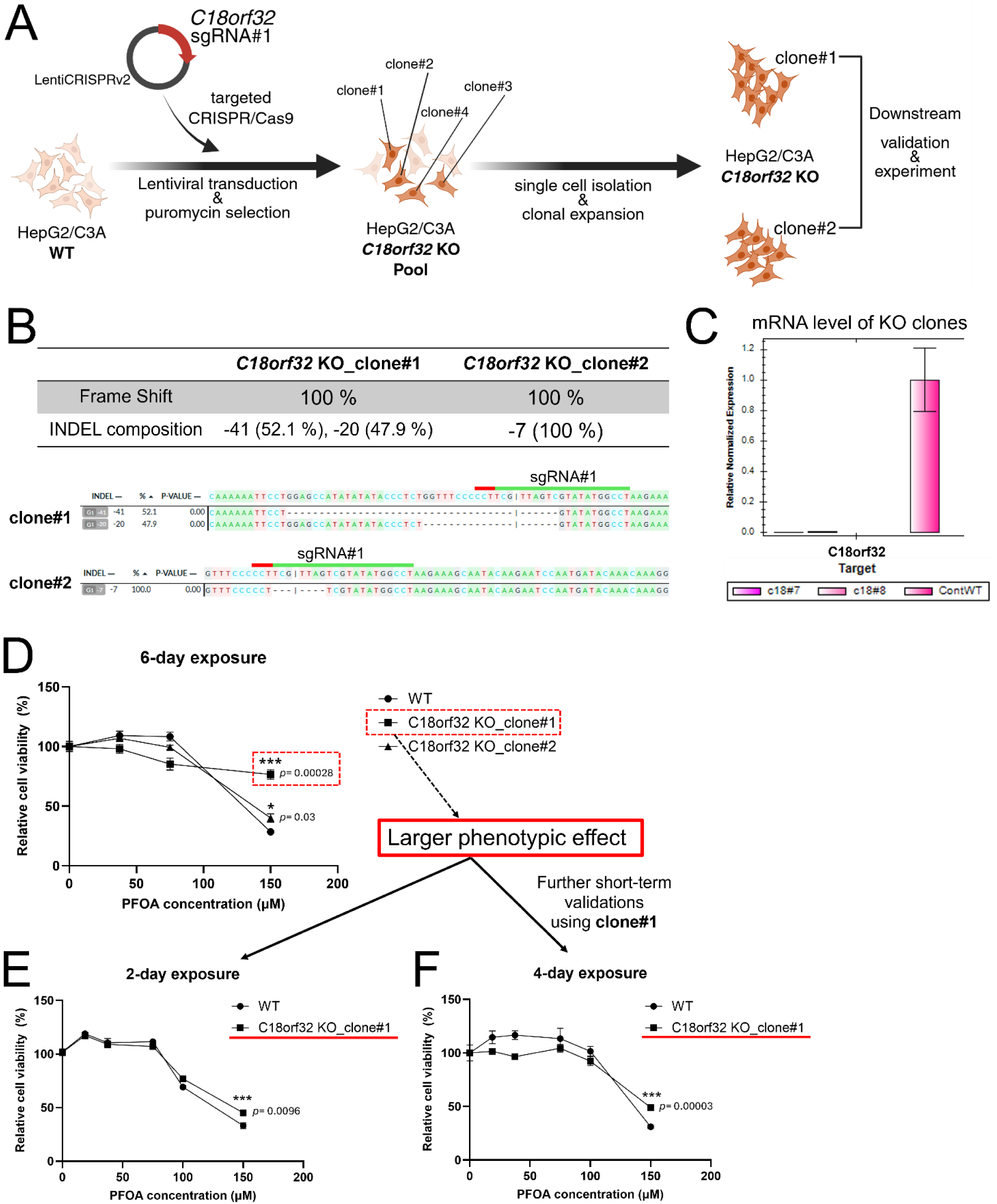
Generation and validation of *C18orf32* knockout HepG2/C3A cell lines. (A) Schematic representation of the CRISPR/Cas9 workflow used to generate *C18orf32* knockout (KO). Wild-type (WT) cells were transduced with LentiCRISPRv2 encoding a *C18orf32*-targeting sgRNA, followed by puromycin selection to obtain a *C18orf32* KO pool. Single-cell clones were isolated and expanded, and representative KO clones (clone#1 and clone#2) were selected for downstream validation and functional studies. (B) Sanger sequencing coupled with DECODR analysis confirms efficient CRISPR-mediated editing at the *C18orf32* locus in selected KO clones. Indel spectra and alignment traces demonstrate high-percentage frameshift mutations disrupting the target site. (C) qRT-PCR analysis shows relative expression level of *C18orf32* transcript in KO clones (clone#1 and clone#2) compared to WT control cells. Bars represent mean ± SD, confirming successful knockout at the mRNA level. (D) Cell viability assays of clone#1 and clone#2 for 6-day exposure of PFOA (0, 32.5, 75, and 150 μM) for phenotypic validation. Shorter-term cell viability validation assays of 2-day (E) and 4-day (F) exposures using clone#1 whose cellular resistance to PFOA is larger than clone#2 in panel (D).

### 3. C18orf32p, a lipid droplet protein, is required for lipid accumulation in liver cells exposed to PFOA

As noted earlier, *C18orf32* (Chromosome 18 open reading frame 32) is a poorly characterized gene encoding a small (76 amino acids), evolutionally conserved, high-confidence LD-associated protein (C18orf32p)^62^. Given recent evidence implicating the functional role of C18orf32p in LD dynamics^62,63^ and prior reports of PFOA-induced lipid metabolic dysregulation^16,64^, we evaluated the phenotypic effect of *C18orf32* disruption on LD accumulation in HepG2/C3A cells exposed to PFOA. LD staining using BODIPY 493/503 combined with live-cell fluorescence microscopy revealed that *C18orf32* KO cells exhibited a marked reduction in LD number under basal conditions, with an even more pronounced reduction upon PFOA exposure (100 µM) (Figure 3A). This phenotype suggests that C18orf32p is required for proper LD accumulation and may contribute to PFOA-induced lipid dysregulation. To quantitatively assess neutral lipid storage, we measured triacylglycerol (TAG) levels in WT and *C18orf32* KO cells with or without PFOA exposure using the Triglyceride-Glo™ assay (Figure 3B). Consistent with the imaging results, KO cells displayed less than half the TAG content of WT cells. Furthermore, whereas WT cells exhibited a significant TAG increase following PFOA treatment (1.8-fold increase, p= 0.008), PFOA failed to induce TAG accumulation in *C18orf32* KO cells (Figure 3B). Together, these findings demonstrate that C18orf32p is required for LD and TAG accumulation in human liver cells, particularly under PFOA exposure, supporting its functional role as a critical regulator of PFOA-induced lipid metabolic disruption.

**Figure 3.**
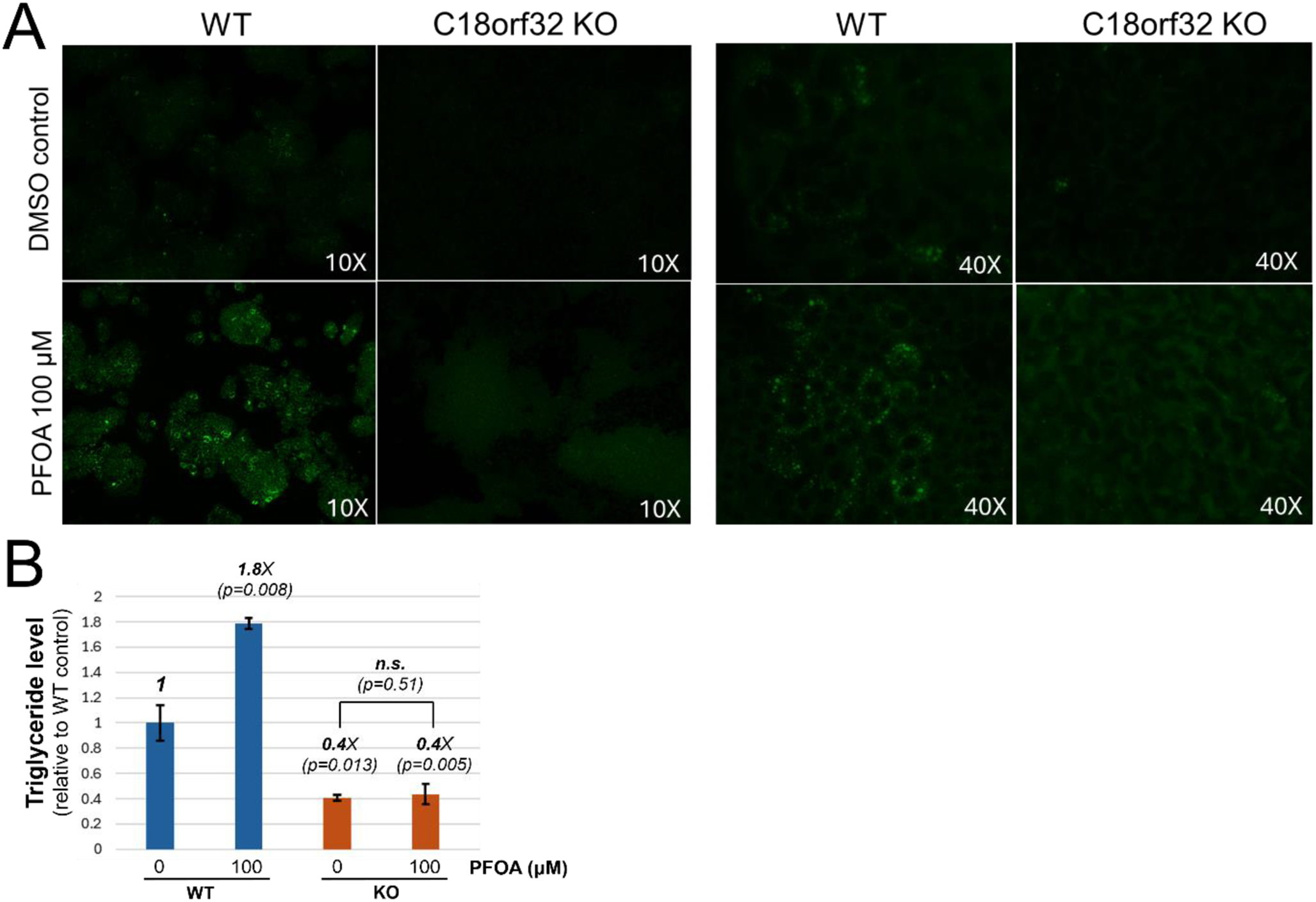
*C18orf32* is required for lipid droplet (LD) and triacylglycerol (TAG) accumulation in human liver cells exposed to PFOA. (A) LD is visualized using BODIPY 493/503 dye (green fluorescence staining for LD) in wile-type (WT), WT+PFOA (100 μM for 72 h), *C18orf32* knockout (KO), *C18orf32* KO+PFOA (100 μM for 72 h). Left-side panels were imaged at 10X magnitude, and right-side panels were imaged at 40X magnitude using the fluorescent microscope. (B) Quantification of TAG levels. A bar graph comparing TAG level (relative to WT control) in WT and *C18orf32* KO cells under control (0 μM) and PFOA exposure (100 μM for 72 h).

### 4. *C18orf32* knockout extensively reprograms the transcriptome of HepG2/C3A liver cells

To investigate the molecular basis underlying the *C18orf32* KO phenotype and its interaction with PFOA-induced toxicity, we profiled the transcriptome of HepG2/C3A cells across four experimental conditions: (1) WT cells, (2) *C18orf32* KO cells, (3) WT cells exposed to PFOA (100 µM, 72h), and (4) *C18orf32* KO cells exposed to PFOA (100 µM, 72h). Principal component analysis (PCA) revealed four distinct transcriptional clusters, with *C18orf32* KO samples showing larger separation from WT than PFOA-treated samples (Figure 4A). Hierarchical clustering also confirmed broad transcriptomic differences between KO and WT cells (Figure 4B), and mRNA abundance of *C18orf32* validated genetic disruption of *C18orf32* (Supplementary Figure S4) which was initially assessed by qRT-PCR (Figure 2C).

**Figure 4.**
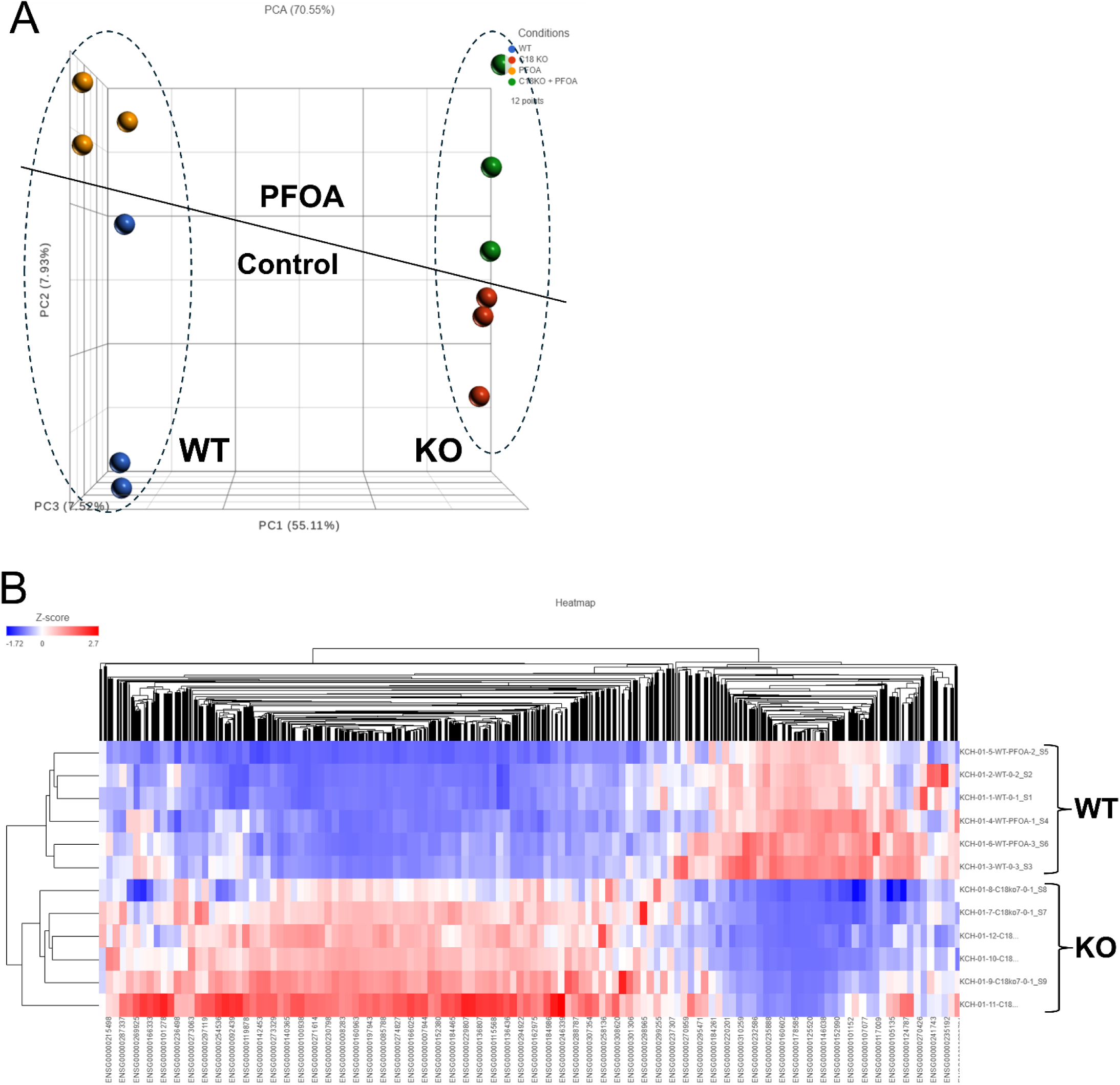
Systematic transcriptome profiling of *C18orf32* KO and WT HepG2/C3A cells in absence and presence of PFOA exposure. (A) Principal Component Analysis (PCA) plot visualizing the overall variance of all experimental conditions in the transcriptomics data. Wild-type (WT)= Blue dots, WT+PFOA: Orange dots, *C18orf32* knockout (KO): Red dots, and *C18orf32*+PFOA: Green dots. (B) Hierarchical clustering heatmap displaying the gene expression levels (z-scores) of a subset of genes across the experimental conditions, with clustering applied to both the genes (rows) and the conditions (columns). The z-score legend shows a scale from blue (low expression, −1.72) to red (high expression, +2.7)

Differential expression analysis identified 3798 significantly increased transcripts (*C18orf32* KO UP set) and 3839 decreased transcripts (*C18orf32* KO DN set) in *C18orf32* KO vs. WT (FDR < 0.01, |log_2_FC| > 2) (Figure 5A, Table S4). GO-BP and KEGG functional enrichment analyses of the corresponding encoded proteins of the *C18orf32* KO DN set revealed marked enrichment for lipid-metabolic processes, including fatty acid β-oxidation, lipid oxidation, fatty acid catabolism, long-chain fatty acid transport, lipid storage, PPAR signaling, cholesterol metabolism, bile secretion, fatty acid degradation, peroxisome pathways, and ferroptosis. Immune-related terms such as complement and coagulation cascades were also enriched, along with multiple amino-acid metabolic pathways (Figure 5B). *The C18orf32* KO UP set is significantly enriched for genes encoding proteins involved in neuronal development, system development, morphogenesis, MAPK signaling, cAMP/Rap1 signaling, cytoskeletal organization, and cardiomyopathy pathways (Figure 5C). In contrast, PFOA exposure alone induced a modest transcriptional response, with only 31 increased (PFOA UP set) and 169 decreased transcripts (PFOA DN set) as compared to unexposed HepG2/CA3 cell (FDR < 0.05, |log_2_FC| > 0.5) (Figure 5D, Table S4). The PFOA DN set is enriched for genes encoding proteins involved in developmental and neuronal processes, cytoskeletal pathways, and axon guidance (Figure 5E). PFOA UP set is enriched in genes encoding proteins in lipid and sterol metabolic processes (sterol biosynthesis, cholesterol homeostasis, triglyceride and acylglycerol homeostasis, VLDL remodeling), as well as ethanol/alcohol metabolism (Figure 5F). KEGG pathways confirmed significant activation of lipid-metabolic programs, including fatty acid metabolism, PPAR signaling, unsaturated fatty acid biosynthesis, fatty acid degradation, peroxisome, and ferroptosis (Figure 5F). Of note, in addition to lipid metabolic pathways, several neuron-related pathways (such as axon guidance) also displayed opposing transcriptional responses between *C18orf32* KO and PFOA exposure. The *C18orf32* KO + PFOA vs. *C18orf32* KO comparison showed only 9 DEGs with no functional enrichment result (Figure 5G) while the *C18orf32* KO + PFOA vs. WT comparison yielded a transcriptome highly similar to the *C18orf32* KO vs. WT, with 4007 upregulated and 3974 downregulated genes, and with functional enrichment patterns closely overlapping those of the *C18orf32* KO vs. WT dataset (Supplementary Figure S5 and Table S4).

**Figure 5.**
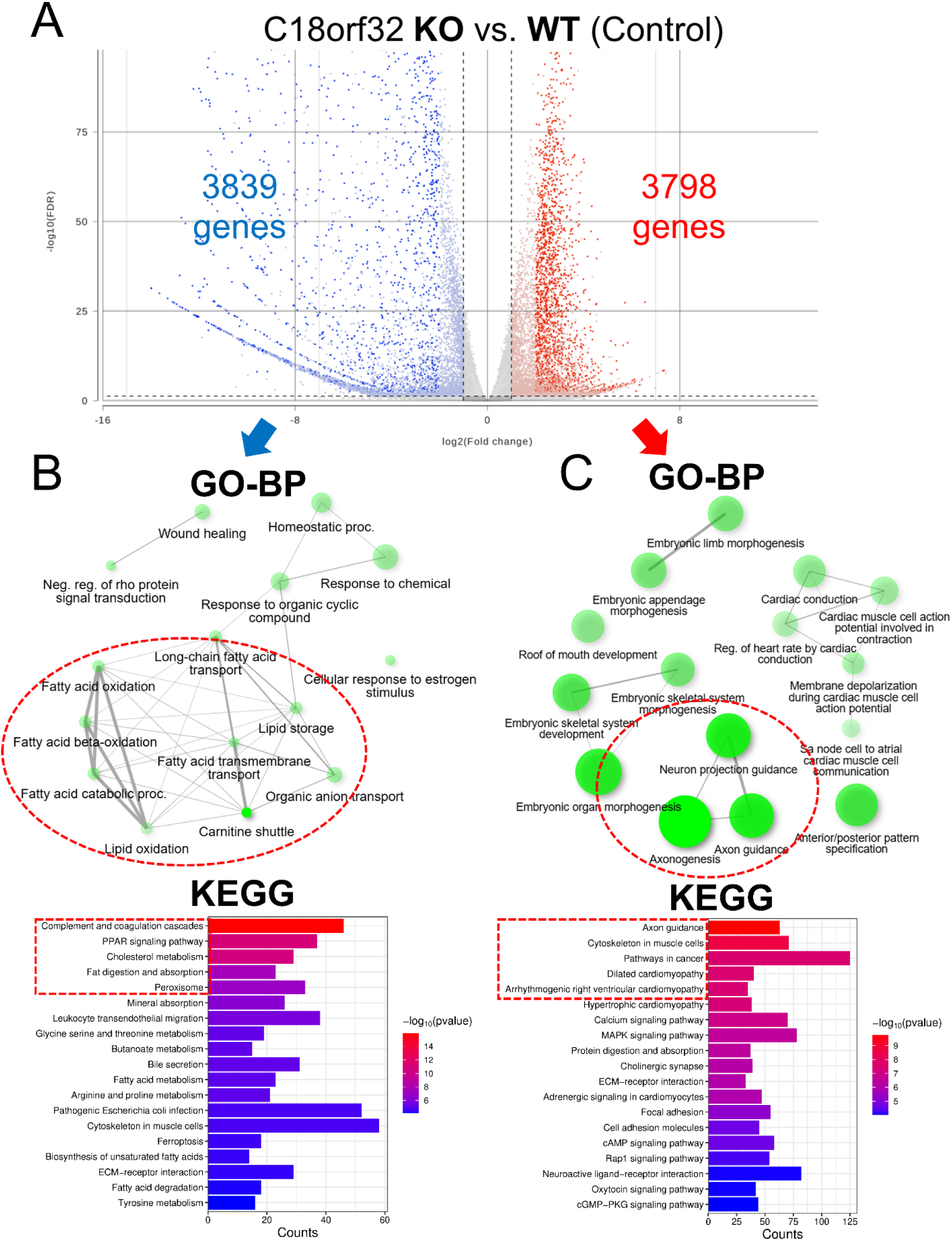
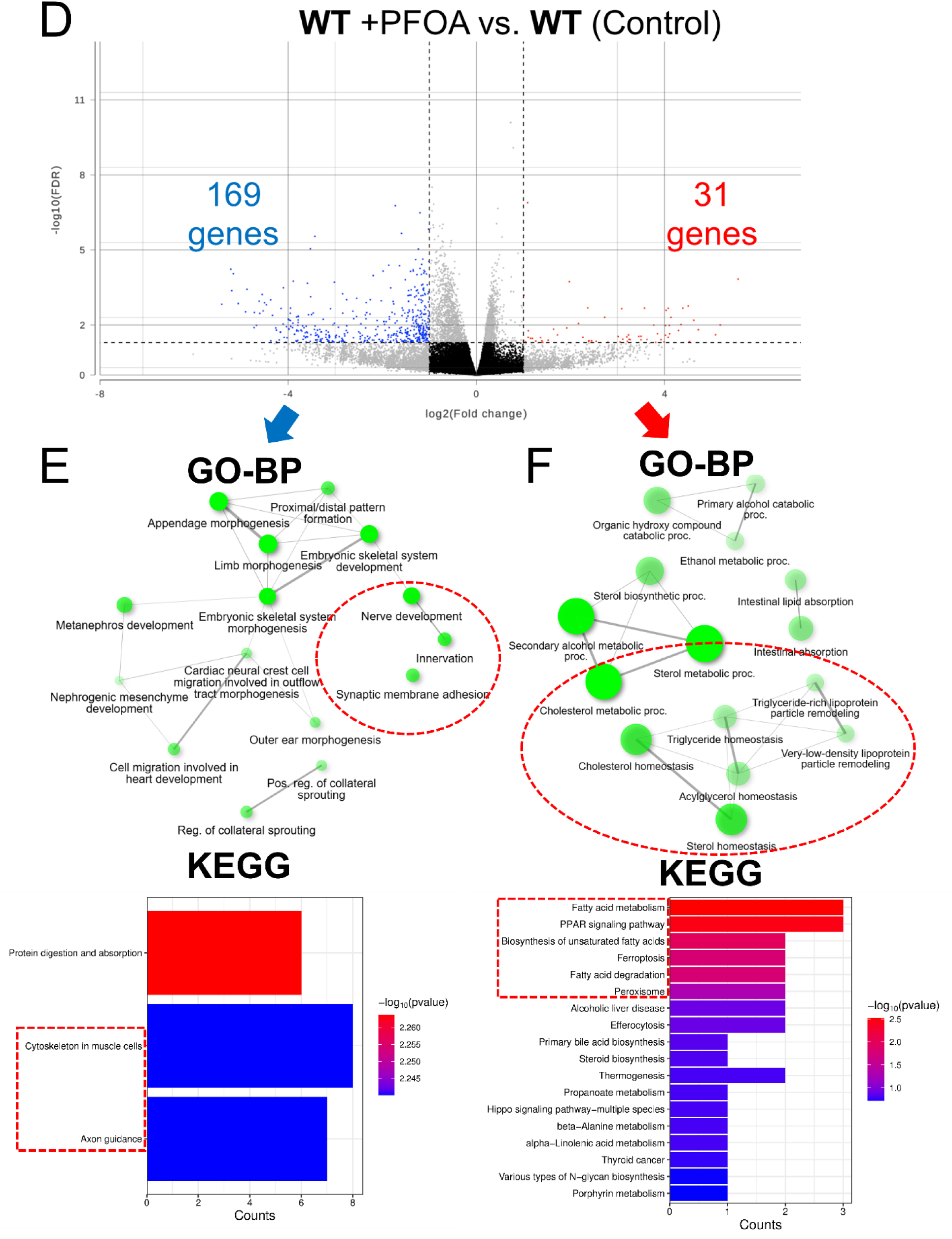
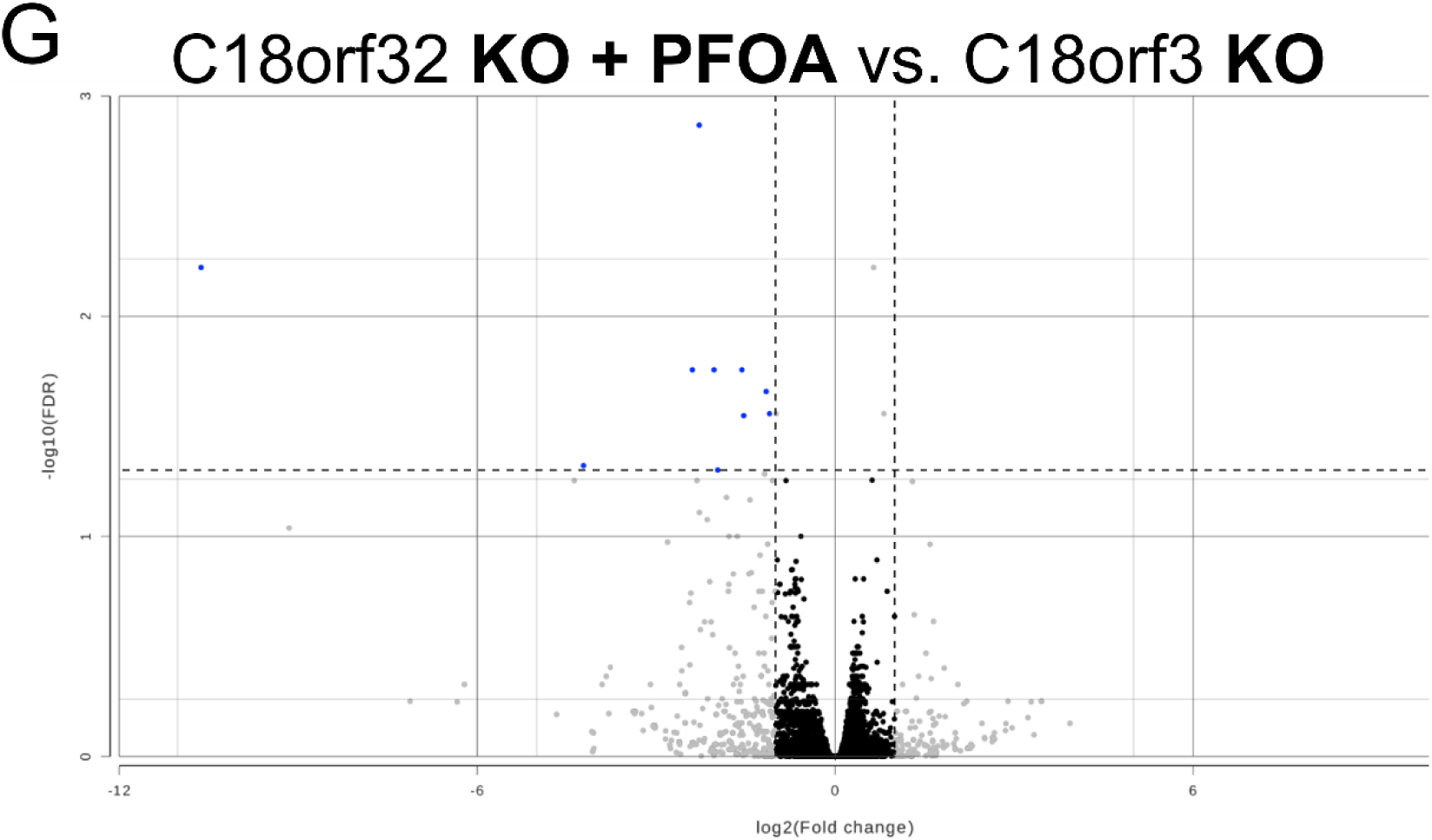
Transcriptomic analysis of *C18orf32* knockout (KO) versus wild-type (WT) and PFOA exposure versus WT (control). (A) Volcano plot visualizing differentially expressed genes (DEGs) of *C18orf32* KO cells compared to WT. Functional pathway enrichment of downregulated genes (B) and upregulated genes (C) using Gene Ontology (GO)-Biological Process (BP) and KEGG. (D) Volcano plot visualizing DEGs of PFOA exposed cells (WT +PFOA) compared to WT (control). Statistically significant downregulated genes (blue dots) and upregulated genes (red dots) are indicated. Functional pathway enrichment of downregulated genes (E) and upregulated genes (F) using Gene Ontology (GO)-Biological Process (BP) and KEGG. (G) Volcano plot visualizing DEGs of *C18orf32* KO cells exposed to PFOA compared to *C18orf32* KO. For A, D, and G, statistically significant downregulated genes (blue dots) and upregulated genes (red dots) are indicated.

### 5. *C18orf32* knockout attenuates PFOA-induced lipid dysregulation by suppressing key lipid metabolism pathways

PFOA has been hypothesized to lead to hepatotoxic effects through activation of PPAR nuclear receptors—particularly PPARα in rodents^20,22^ but less potent in humans^65,66^—which has been suggested as a central mechanism underlying its toxicity. Given the contrasting transcriptomic responses observed between *C18orf32* KO, WT, PFOA-exposed *C18orf32* KO, and PFOA-exposed WT cells—in particular the divergent expression of lipid metabolic pathways, we performed a targeted analysis of key lipid metabolism pathways to identify potential pathway-level changes. We selected 8 representative lipid metabolic pathways, focusing on those showing the most striking differences, and examined the mRNA expression patterns of individual constituent genes (Figure 6).

**Figure 6.**
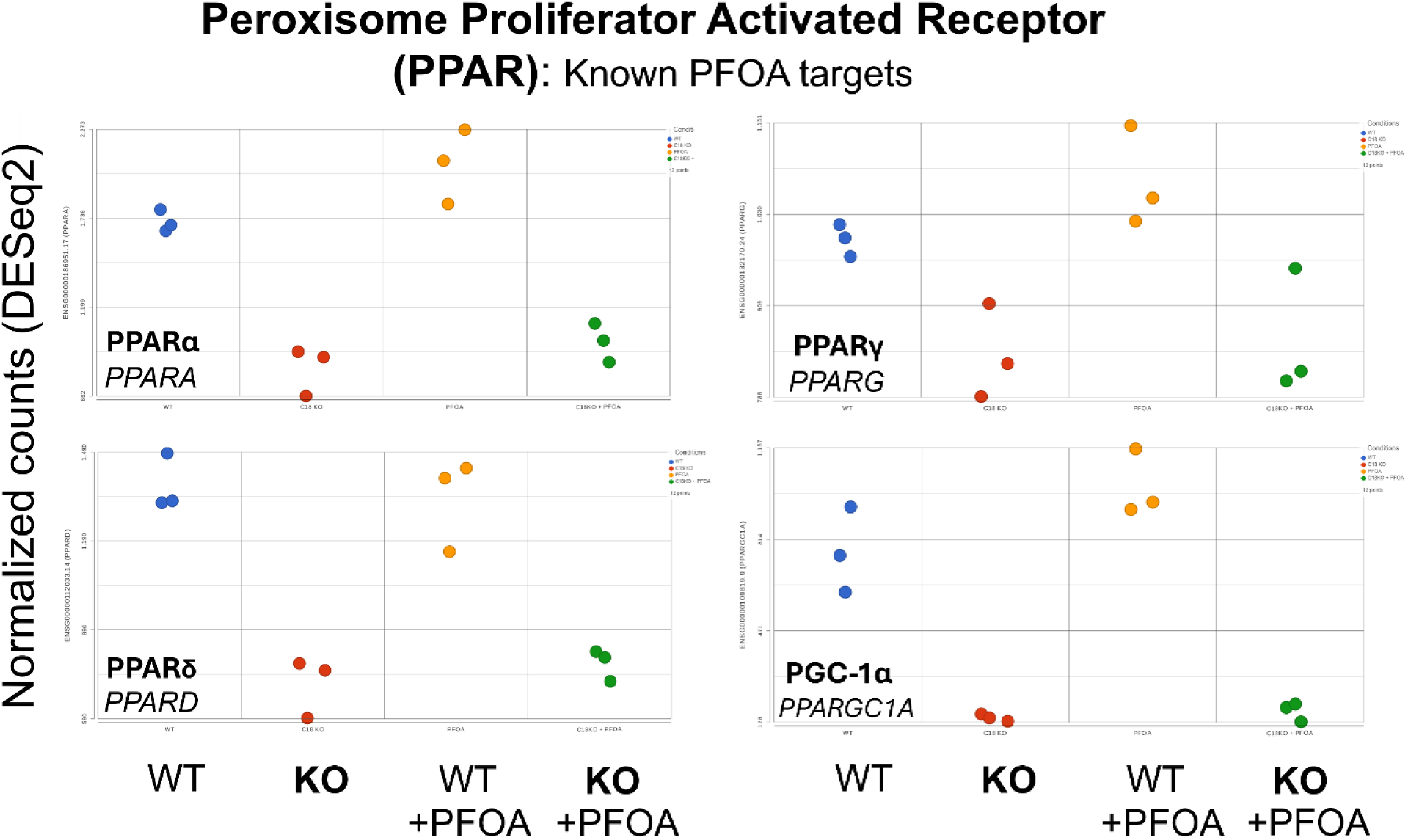
*C18orf32* knockout (**KO**) impact on peroxisome proliferator activated receptors (PPAR) and a key coactivator (*PPARA*, *PPARG*, *PPARD*, and *PPARGC1A*). Individual mRNA expression plots showing the normalized counts (y-axis) of a specific gene across the four experimental conditions (x-axis): WT, *C18orf32* KO, WT + PFOA, and *C18orf32* KO + PFOA.

**Figure 7.**
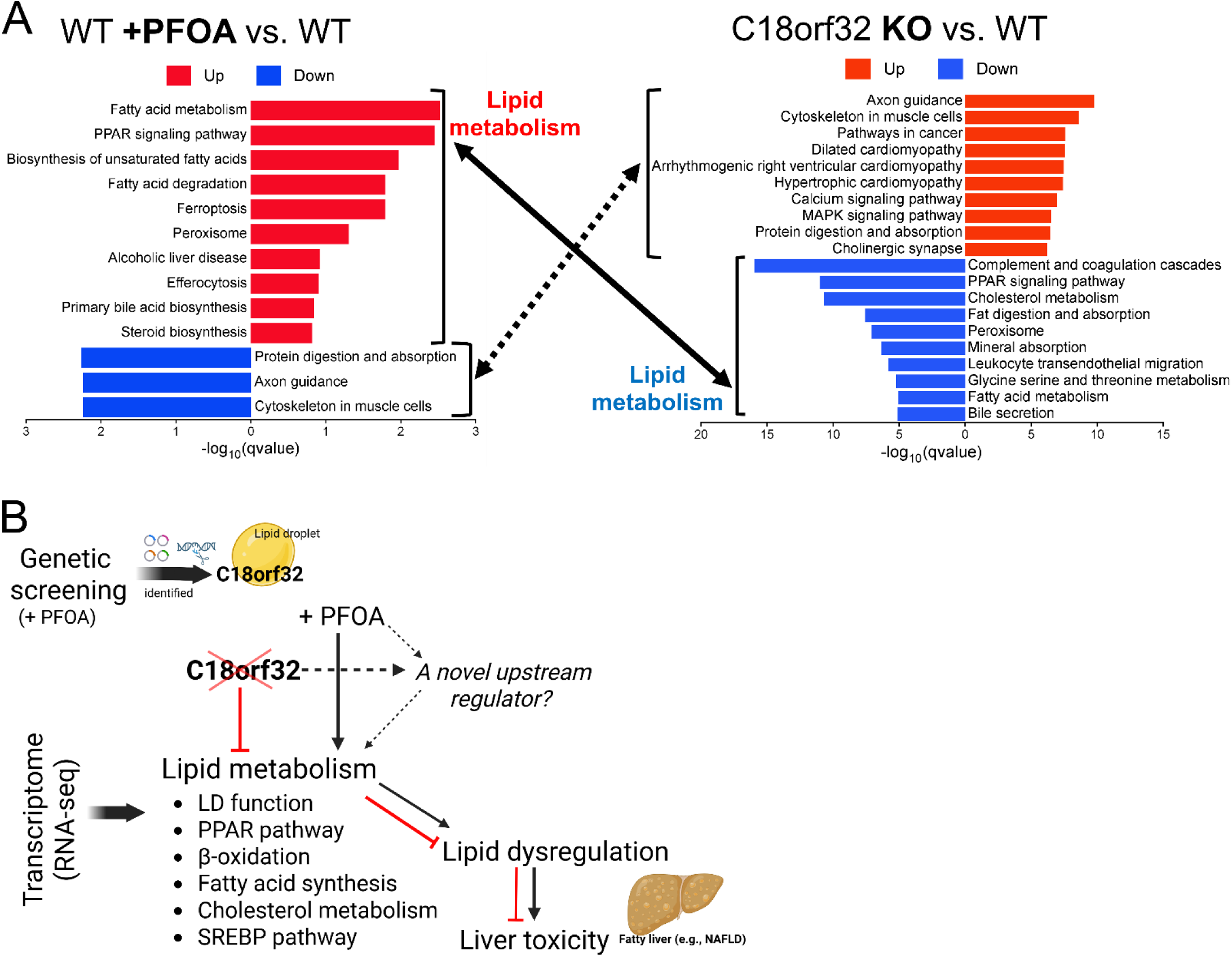
Proposed molecular mechanism linking C18orf32p to lipid metabolic regulation during PFOA exposure. (A) Overview of the core mechanistic insight (using top 10 KEGG pathways) showing that loss of *C18orf32* desensitizes cells to PFOA-induced toxicity by attenuating the disruption of lipid metabolic pathways normally triggered by PFOA exposure. (B) Schematic summary illustrating C18orf32p as a novel upstream regulator of lipid metabolism that contributes to the lipid dysregulation underlying PFOA-induced hepatotoxicity.

Expression of core components of the PPAR signaling pathway—*PPARA*, *PPARG*, *PPARD*, and *PPARGC1A*—were markedly reduced in *C18orf32* KO cells under basal conditions compared to WT cells. In WT cells, PFOA exposure modestly increased mRNA levels of PPAR family members and *PPARGC1A* (except for *PPARD*); however, this response was significantly attenuated or absent in *C18orf32* KO cells (Figure 6 and Table 1). Similarly, mRNA levels of nuclear receptors implicated in PPAR-independent PFAS toxicity, including *PXR, FXR, RXR,* and *CAR*, were substantially reduced in the absence of *C18orf32* and remained low following PFOA exposure (Table 1). We next examined genes encoding Acyl-CoA oxidases (*ACOX1, ACOX2,* and *ACOX3*), key enzymes in peroxisomal β-oxidation and canonical PPARα downstream targets. Expression of all three *ACOX* genes was significantly lower in *C18orf32* KO cells relative to WT cells under both basal and PFOA-treated conditions (Table 1). Likewise, mRNA levels of Perilipin (PLIN) family members (*PLIN2, PLIN3,* and *PLIN4*), which encode structural components essential for lipid droplet formation and stability, were markedly reduced in KO cells, with PFOA-induced upregulation observed in WT cells but not in KO cells (Table 1). We further evaluated sterol regulatory element-binding proteins (SREBPs), master transcriptional regulators of fatty acid, triglyceride, and cholesterol biosynthesis. mRNA expression levels of both *SREBF1* and *SREBF2* were significantly decreased in *C18orf32* KO cells (Table 1). Consistent with reduced SREBF1/2 expression, multiple downstream metabolic enzymes exhibited parallel decreases in transcript abundance, including de novo fatty acid synthesis enzymes (*ACLY, ACSS3, FASN*), triacylglycerol synthesis enzymes (*DGAT2, ACSL3*), and cholesterol biosynthesis and esterification enzymes (*LSS, SQLE, SOAT1, and FDFT1*) (Table 1).

**Table 1.**
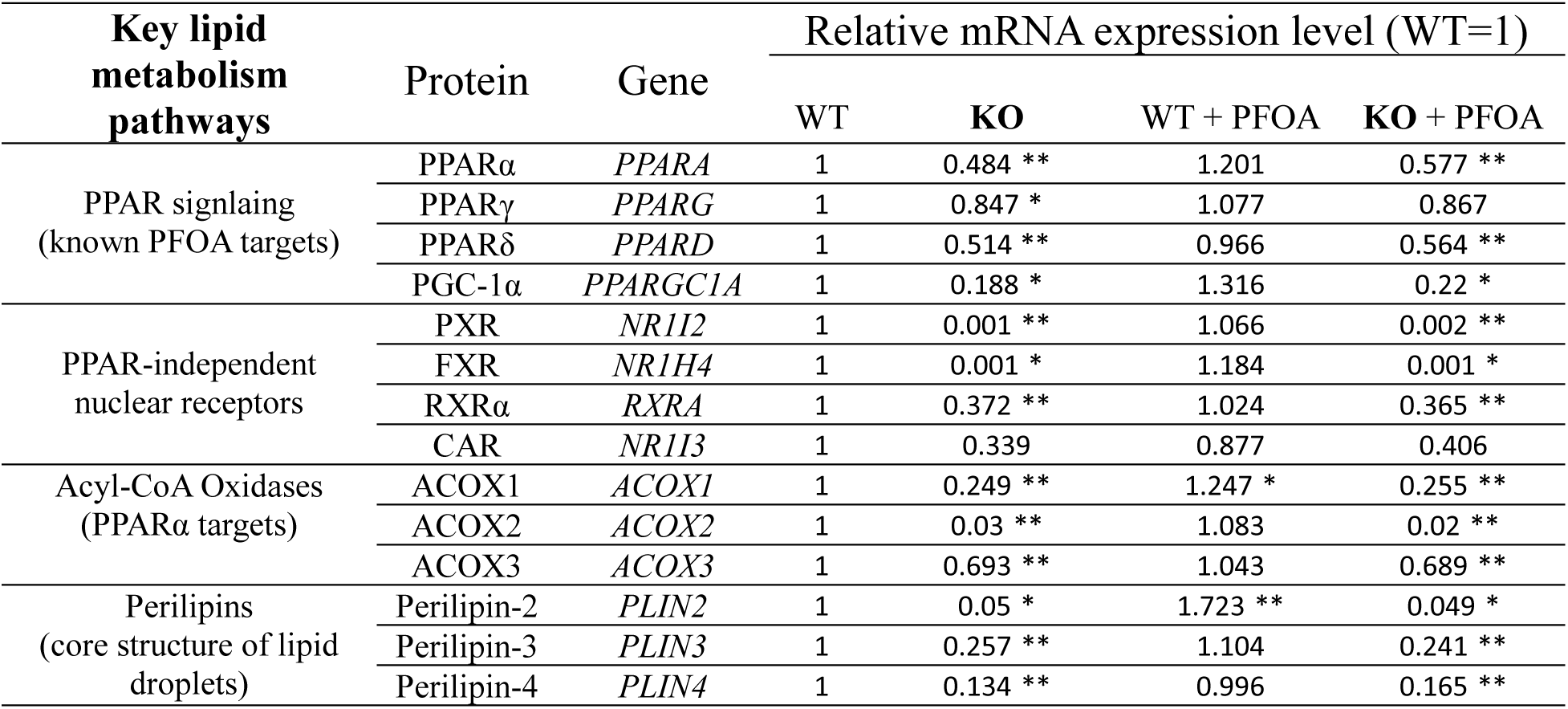
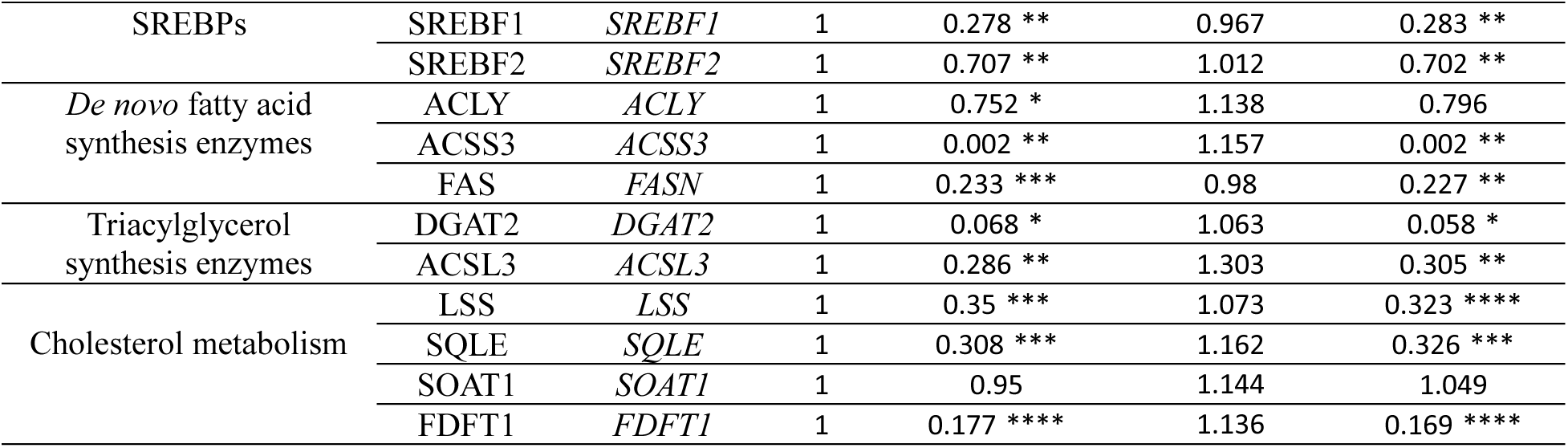
*C18orf32* knockout (KO) impact on gene components of key lipid metabolism pathways. Relative mRNA expression level of a specific gene across the four experimental conditions: Wild-type (WT), *C18orf32* KO, WT + PFOA, and *C18orf32* KO + PFOA. RNA-seq normalized counts were used to calculate the relative level followed by two-sided t-test for statistical significance. *p < 0.05, **p < 0.01, ***p < 0.001, ****p < 0.0001. PPAR, peroxisome proliferator activated receptor; SREBPs, sterol regulatory element-binding proteins.

## Discussion

Despite extensive research, the molecular basis underlying human-relevant PFOA—and PFAS more broadly—induced hepatotoxicity and lipid dysregulation remain incompletely understood. Here, we employed a functional toxicogenomics strategy to further understand PFOA-induced human liver toxicity. Our genome-wide CRISPR KO screen identified multiple candidate genes whose loss either sensitized or protected cells from PFOA-induced cytotoxicity. The candidates were enriched for lipid metabolism and energy-related pathways, including fatty acid metabolism, oxidative phosphorylation, peroxisomal function, and cholesterol homeostasis. Among these candidate genes, we focused on *C18orf32* for downstream mechanistic analyses based on two key considerations: (1) its robust and reproducible high-ranking performance across different bioinformatic scoring algorithms, and (2) its established connection to lipid metabolism as a critical component of LDs. Previous studies^62,67^ reported changes in lipid species and LD accumulation/secretion patterns due to partial disruption of *C18orf32* but the functional role of C18orf32p in toxicant response was previously uncharacterized. LDs are present in most organisms and mammalian tissues^63,68^ and C18orf32p has orthologs across vertebrate and invertebrate species ([Unknome v3](https://unknome.mrc-lmb.cam.ac.uk/cluster_details/UKP09357/0/?utm_source=chatgpt.com)). BLAST^69^ and DIOPT Orthology ([DRSC – DRSC Integrative Ortholog Prediction Tool](https://www.flyrnai.org/cgi-bin/DRSC_orthologs.pl)) analyses of human C18orf32p illustrated evolutionary conservation across metazoans (Supplementary Figure 6 and Table S5). In people, predicted loss-of-function mutations in *C18orf32* results in an autosomal recessive neurodevelopmental disorder, characterized by global developmental delay, structural brain anomalies, and hypotonia^70^. Given the conservation of both LD biology and C18orf32p, PFAS-induced lipid dysregulation mechanisms observed in our study may have one-health relevance^71,72^.

### *C18orf32* knockout establishes a key role in regulating PFOA cytotoxicity and lipid accumulation

Targeted CRISPR/Cas9 KO of *C18orf32* validated the screening phenotype. Although the two KO clones exhibited different degrees of resistance, *C18orf32*-deficient hepatocytes exhibited enhanced resistance to PFOA-induced cytotoxicity and showed marked reductions in LD number and TAG abundance, particularly under PFOA exposure. These findings suggest that loss of *C18orf32* diminishes LD availability and TAG storage, thereby attenuating PFOA-induced lipid dysregulation and conferring functional resistance. This phenotype aligns with it classification as a high-confidence core LD protein^62^ and with reports of PFOA-induced LD accumulation in *C.elegans*, mouse hepatocytes, bovine blastocytes, and human HepaRG cells^31–35^. PFOA-induced neutral lipid accumulation has been linked to diverse adaptive and maladaptive lipogenesis responses^16,30,73,74^. We found significantly decreased mRNA levels of genes involved in those pathways in *C18orf32* knockout alone or after exposure to PFOA (Figure 6) and thus supports a role for C18orf32p as a critical modulator of TAG synthesis, LD biogenesis, or LD maturation^63,75^ after PFOA exposure. Importantly, our results support a functional linkage between*C18orf32*-dependent LD accumulation and PFOA-induced hepatotoxicity, establishing a mechanistic bridge between PFOA exposure, lipid metabolism dysregulation, and toxicity outcomes. And more broadly, if C18orf32p contributes to human-relevant lipid accumulation responses to PFAS in general or those related to PFOA, it may help explain PFAS-induced steatosis phenotypes.

### *C18orf32* loss triggers extensive transcriptomic reprogramming of lipid metabolic networks

Transcriptomic profiling revealed that *C18orf32* KO elicits more profound global expression changes than PFOA exposure alone, with an unexpectedly large number of DEGs. This widespread transcriptomic reprogramming indicates that C18orf32p likely functions as a key regulator of essential metabolic processes, particularly those governing lipid metabolism^76,77^. Consistent with this interpretation, mRNA levels of genes encoding proteins involved in major lipid pathways, including fatty acid synthesis, TAG synthesis, cholesterol biosynthesis, peroxisomal β-oxidation, and PPAR signaling—were markedly decreased in *C18orf32*-deficient cells as compared to WT cells. Individual gene-expression analyses further demonstrated the regulatory patterns of these key lipid-metabolism pathways. In particular, pathways such as cholesterol synthesis and fatty-acid metabolism—largely governed by SREBP transcription factors—were universally lower in *C18orf32*-deficient cells. SREBP1 (*SREBF1*) drives fatty-acid and TAG synthesis and promotes LD formation, whereas SREBP2 (*SREBF2*) regulates cholesterol biosynthesis and LDL uptake^78,79^. These SREBP-dependent lipogenic and cholesterologenic pathways are frequently activated by PFAS exposure^80–82^ and contribute to human-relevant steatosis and lipid dysregulation^82^. Human hepatocytes exhibit relatively weak PPARα response to PFOA and other PFAS, and human-relevant toxicity has been associated with alterations in PPARγ, PPARδ, CAR, PXR, FXR, and RXR signaling, which regulate lipid metabolism, bile-acid homeostasis, and xenobiotic responses^16,23,39,83,84^. mRNA levels of all these nuclear receptors were suppressed in *C18orf32*-deficient cells, suggesting that *C18orf32* deficiency impacts multiple signaling pathways associated with PFOA toxicity mechanism in addition to PPARα. These findings reinforce the hypothesis that PPARα is not the dominant driver of PFOA toxicity in human cells and suggests alternative mechanisms underly the biological effects of PFOA, with C18orf32p as a novel molecular target of PFOA toxicity and lipid dysregulation.

We also observed strong suppression of *PLIN2, PLIN3,* and *PLIN4* in *C18orf32* KO cells. PLIN proteins form the structural LD coat, stabilize LDs, and mediate lipid-signaling pathways that promote neutral-lipid storage^85,86,87^. The coordinated suppression of mRNA expression highlights the functional dependence of LDs on C18orf32p. LDs are now recognized as dynamic organelles that integrate lipid flux, organelle communication, proteostasis, and stress-response signaling^88,89^. Through interactions with mitochondria, peroxisomes, lysosomes, and the ER by coordinating β-oxidation, fatty-acid trafficking, membrane lipid synthesis, and redox balance LDs serve as central hubs that maintain lipid metabolic and transcriptional homeostasis^63,88,90,91^. Emerging evidence also implicates LDs in the sequestration of potentially harmful oxidized lipids and the modulation of reactive oxygen species (ROS) accumulation^63,92^. These functions suggest that LDs not only buffer lipid overload but also protect cells from oxidative stress induced by excess lipids, a mechanism that may contribute to the resistant phenotype observed in our *C18orf32* KO models.

and protect cells from oxidative stress induced by excess lipid load. Thus, disruption of a core LD component such as C18orf32p may produce broad transcriptomic consequences. In this context, the extensive transcriptomic reprogramming observed in *C18orf32*-deficient liver cells—dominated by lipid-metabolism pathways—demonstrates how tightly LD integrity is embedded within cellular metabolic architecture. Collectively, these findings suggest that *C18orf32* KO rewires the hepatocyte lipid-metabolic network in a manner that not only desensitizes the cell to PFOA-induced lipid dysregulation and hepatotoxicity but may also reshape cellular responses to other PFAS or lipid-active toxicants. This reprogramming provides a mechanistic explanation for the observed resistance to PFOA cytotoxicity, including reduced LD abundance and lower TAG levels in *C18orf32*-deficient cells.

### Mechanistic and translational implications

Given its strong and broad regulatory influence over lipid metabolic programs, C18orf32p may also serve as a biomarker of LD dysfunction or a susceptibility factor for PFOA-induced liver injury—as well as for structurally related PFAS, including PFBA, PFHxA, GenX, and PFDA. These findings also suggest that targeting C18orf32p or LD regulatory pathways could modulate susceptibility to PFAS-induced hepatotoxicity, offering potential translational opportunities for therapeutic or preventive strategies for liver disease, such as non-alcoholic fatty liver disease (NAFLD)^93,94^. More broadly, this functional genomic framework can be readily applied to other PFAS homologs or environmental chemicals that disrupt lipid metabolism, providing a scalable blueprint for systematic, human-relevant mechanistic toxicology.

### Limitations and Summary

There are several limitations in the study. First, the study relied primarily on HepG2/C3A cells in the genome-wide CRISPR screening and these cells do not fully recapitulate primary human hepatocyte physiology^95–97^. Second, we used high dose relatively short-term exposures in our studies rather than chronic, low-dose exposure conditions that better represent environmental PFAS exposures^81,98^. In addition, our primary endpoint was cellular toxicity as assessed by cellular survival and proliferation in the presence of PFOA and may miss other functionally relevant cellular endpoints. Despite these limitations, this study identified C18orf32p as a critical LD-associated modulator of human hepatocyte responses to PFOA, linking LD formation, TAG accumulation, and transcriptional remodeling of lipid metabolism pathways. By uncovering a PPARα-independent, human-relevant mechanism of PFAS hepatotoxicity, our findings provide new mechanistic insight, validate the power of functional toxicogenomics, and establish a broadly applicable framework for elucidating molecular pathways underlying environmental chemical toxicity.

## Notes

### Competing Interest Statement

The authors have declared no competing interest.

### Funding

- National Institute of Environmental Health Sciences: R01ES033625

