## Supplementary Figures for "Genome-wide CRISPR screens identified *C18orf32* as a novel regulator of lipid metabolism that mediates PFOA-induced toxicity"

*Supplementary Figures S1 to S6.*

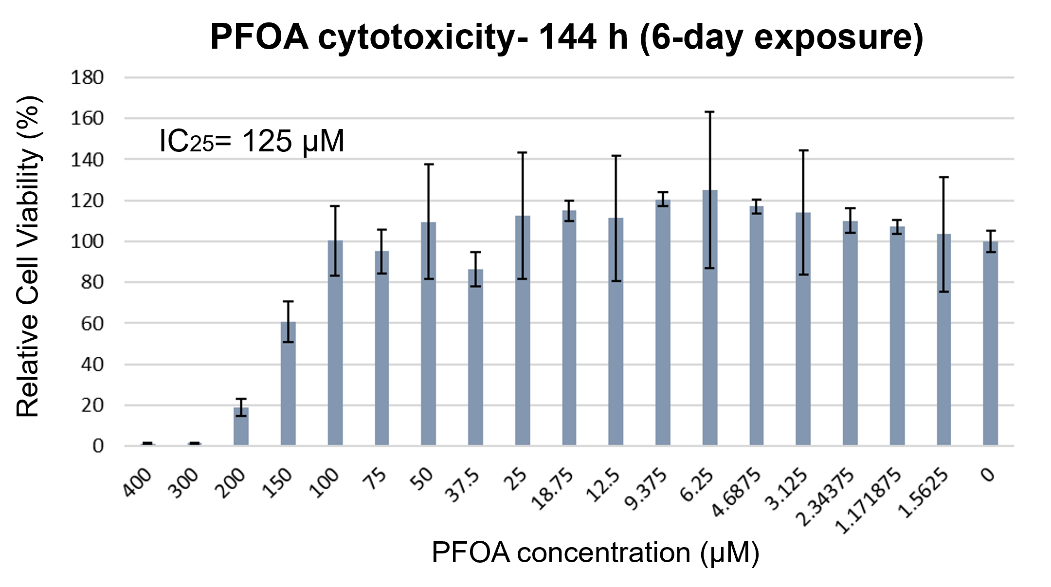

**Figure S1**. PFOA cytotoxicity assessment to determine IC_25_ concentration of PFOA for 144 h (6 days exposure). Cell viability of HepG2/C3A cells exposed to PFOA was assessed in a dose-response manner. The IC_25_ (inhibitory concentration at which the chemical reduces cell viability by 25%) value was calculated using the dose-response sigmoid function in GraphPad Prism. The IC_25_ of PFOA was 125 µM for 6-day exposure.

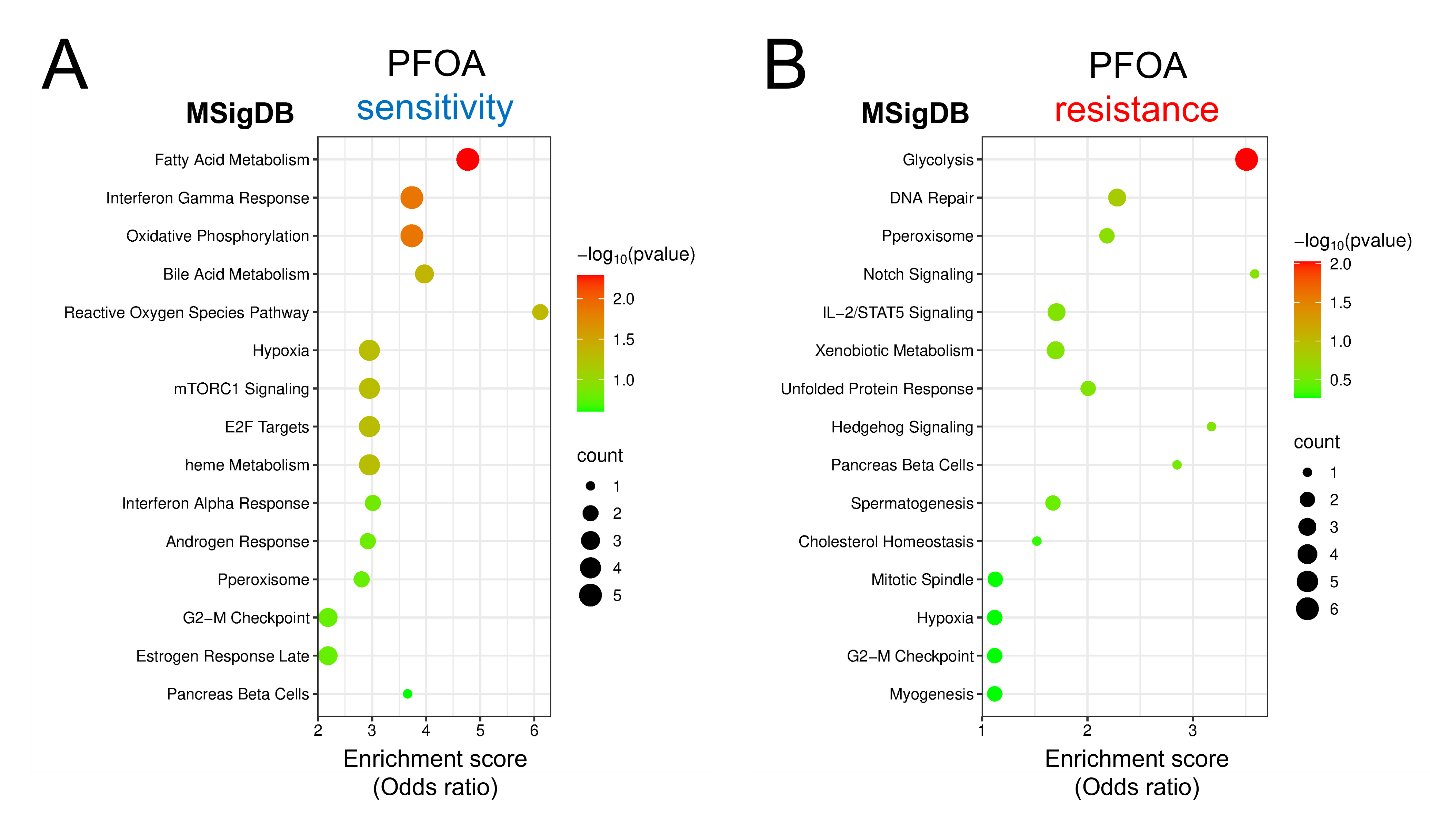

**Figure S2**. Functional enrichment results of PFOA-sensitive and resistant genes using [Enrichr](https://maayanlab.cloud/Enrichr/). Top 15 MSigDB terms are presented for each gene set.

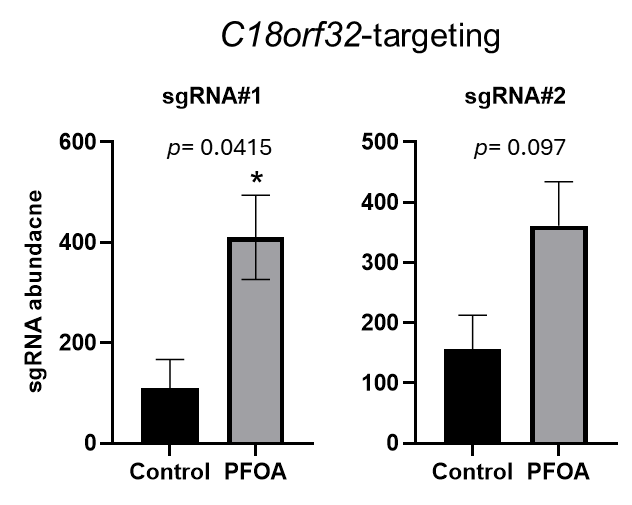

**Figure S3**. Individual sgRNA abundance profiles for *C18orf32* (sgRNA#1 and sgRNA#2) under control and PFOA exposure from the CRISPR screen.

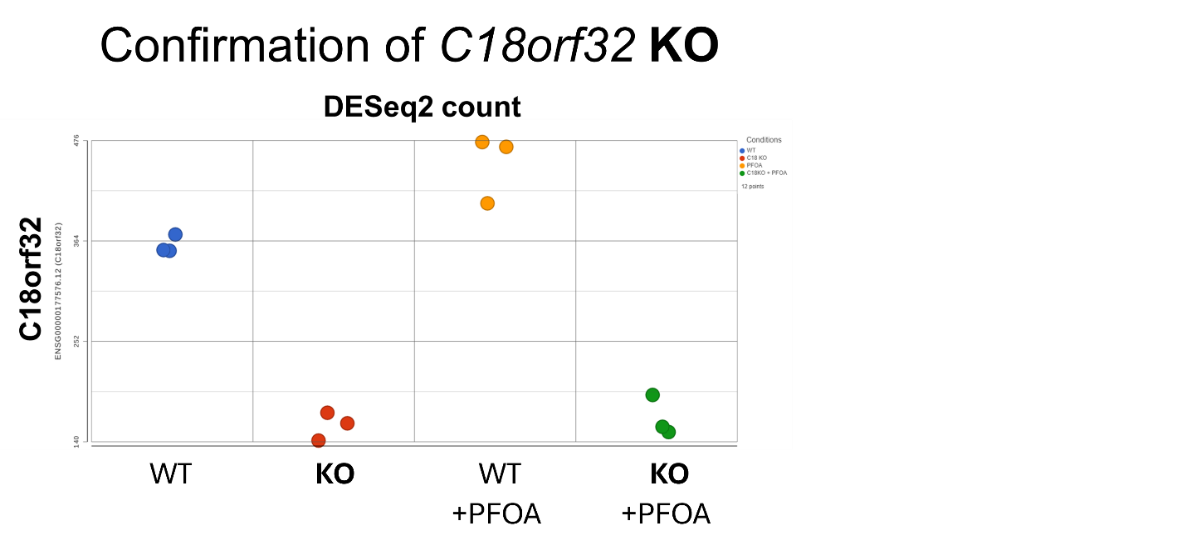

**Figure S4**. *C18orf32* expression plot for KO confirmation which shows the expression level of the *C18orf32* gene across the four experimental conditions. The x-axis represents the four experimental conditions, and the y-axis indicates the expression level of *C18orf32* gene.

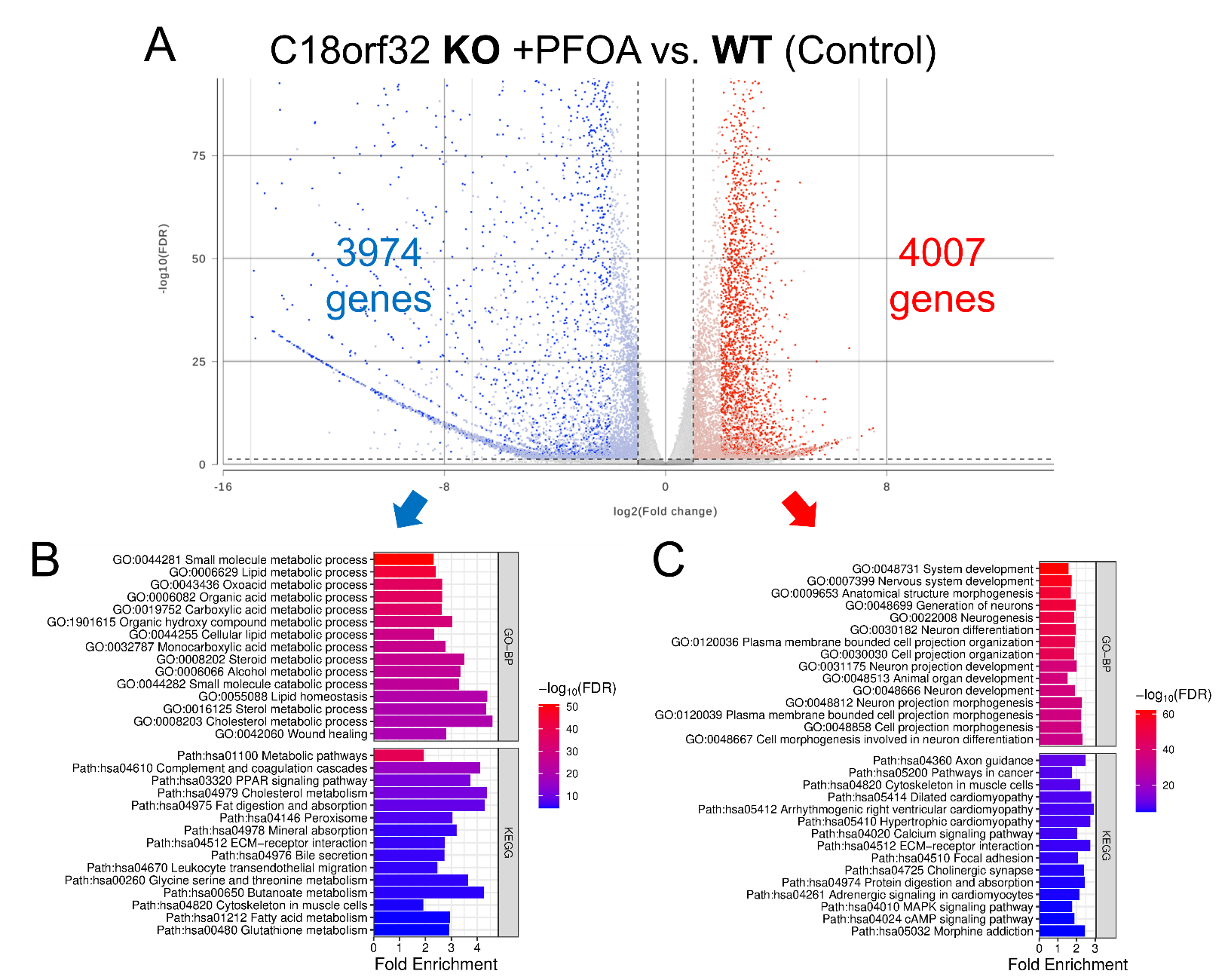

**Figure S5**. Transcriptomic analysis of C18orf32 knockout (KO) cells exposed to PFOA (100 μM for 72 h) versus wild-type (WT) cells. (A) Volcano plot visualizing differentially expressed genes (DEGs). Statistically significant downregulated genes (blue dots) and upregulated genes (red dots) are indicated. Functional pathway enrichment of downregulated genes (B) and upregulated genes (C) using Gene Ontology (GO)-Biological Process (BP) and KEGG.

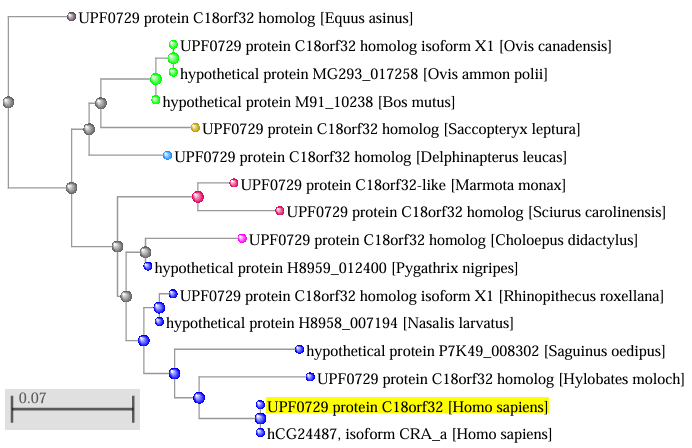

**Figure S6**. BLAST analysis of human C18orf32p. Distance tree of the result is displayed. Scale bar at the bottom left indicates the genetic distance where shorter branches mean higher similarity.
